# Structural models of mitochondrial uncoupling proteins obtained in DPC micelles are not physiologically relevant for their uncoupling activity

**DOI:** 10.1101/2020.07.20.195602

**Authors:** Mathilde S. Piel, Sandrine Masscheleyn, Frédéric Bouillaud, Karine Moncoq, Bruno Miroux

## Abstract

Uncoupling protein 1 (UCP1) is found in the inner mitochondrial membrane of brown adipocyte. In the presence of long-chain fatty acids (LCFA), UCP1 increases the proton conductance, which, in turn, increases fatty acid oxidation and energy release as heat. Several atomic models of UCP1 and UCP2 have been obtained by NMR in dodecylphosphocholine (DPC), a detergent known to inactivate UCP1. Based on NMR titration experiment on UCP1 with LCFA, it has been proposed that K56 and K269 are crucial for LCFA binding and UCP1 activation. Given the numerous controversies on the use of DPC for structure-function analyses of membrane proteins, we revisited those UCP1 mutants in a more physiological context by expressing them in the mitochondria of *S. cerevisiae*. Mitochondrial respiration, assayed on permeabilized spheroplasts, enables the determination of UCP1 activation and inhibition. The K56S, K269S and K56S/K269S mutants did not display any default in activation, which shows that the NMR experiments in DPC detergent are not relevant to understand UCP1 function.

## Introduction

Uncoupling protein 1 belongs to the mitochondrial carrier family^1^. In contrast to other members of the family, which transport anions substrates such as the ADP, ATP, substrates of the Krebs cycle and other catabolic pathways, UCP1 acts as a passive proton transporter. UCP1 is almost exclusively present in Brown Adipose Tissues (BAT) and its function is to ensure non-shivering thermogenesis upon cold exposure^2^. The *Ucp1* gene is highly expressed in BAT and the corresponding protein accumulates at up to 10% of inner mitochondrial proteins^3^. Consequently, the proton permeability induced by UCP1 activity enables to fully uncouple respiration from ATP synthesis and to dissipate energy from oxidative catabolism as heat^4^. BAT is present in all mammals at birth and has been recently rediscovered in some adult humans^5,6^. Given the impact of BAT activity in heat production and body weight control in small mammals, it has been hypothesized that modulating UCP1 activity in human BAT could impact body weight and increase resistance toward obesity and diabetes^7,8^. UCP1 is activated by free fatty acids (FFA) and inhibited by purine nucleotides^9^. However, at molecular levels, the mechanism of proton transport remains elusive and the relationship between FFA and proton translocation is controversial. In the absence of high resolution structural information, several models have been proposed: i) an allosteric model in which binding of fatty acids triggers conformational changes and H^+^ or OH^-^ translocation^4,9,10^; ii) a “H^+^ buffering” model in which carboxylic group of fatty acids participates directly to the H^+^ translocation pathway^11^; iii) a “fatty acid cycling” model, in which UCP1 is a fatty acid anion exporter that stimulates protonation of FFA in the intermembrane space, the passive diffusion of protonated FFA through the inner membrane and consequently the release of proton in the matrix side^12^; iv) a consensus model based on direct measurement of proton current on BAT mitochondria, which proposes the FFA cycling model for short chain FFA and the buffering model for long chain FFA.^13^

The mitochondrial carriers derive from the triplication of a gene encoding a peptide of 100 amino-acids containing two membrane spans.^14,15^ Consequently, UCP1 consist of six transmembrane α-helices, both C-ter and N-terminal ends facing the inter mitochondrial membrane space.^16^ Atomic models obtained from X-ray diffraction of 3D crystals of the bovine and yeast ADP/ATP carrier (AAC) confirmed the pseudo three-fold symmetry for all mitochondrial carriers and revealed an alternating access mechanism of transport driven by two highly conserved salt bridge networks on the matrix and cytoplasmic sides of central cavity.^17–19^ So far all structural data obtained by 3D crystallization required the presence of nanomolar affinity inhibitors blocking the protein in the matrix or cytoplasmic conformational state and no high-resolution structure of mitochondrial carriers have been obtained in presence of their natural substrate. Therefore and despite the prediction sequence analysis of a common substrates binding site within the central cavity^20^, little is known about the substrate translocation pathway. To address the dynamic of mitochondrial carriers, Chou and colleagues used liquid state NMR. Berardi *et al*.^21^ produced UCP2, a mitochondrial carrier sharing 60% sequence identity with UCP1^1^, in *E. coli*, refolded the protein in presence of dodecylphosphocholine detergent (DPC) also called Foscholine-12 and used several innovative NMR and modeling technics to build a structural model of UCP2. In Zhao *et al*.^*22*^ they followed up the initial work on UCP2 with UCP1, with the objective to identify amino-acids involved in FFA binding. Authors refolded UCP1 in DPC, used fatty acids having 16 carbons (C16FA) as activator of the protein to perform chemical-shift titration on purified UCP1 and identified several residues showing binding saturation with apparent K_D_ of 700 +/-200 μM. They concluded that the C16FA binds at the H1/H6 interface and that the driving force of FA binding is mediated by electrostatic interactions between the acidic function of FA and UCP1 basic residues K56 and K269. Reconstitution of the purified UCP1 in proteoliposomes seemed to confirm that proton transport activity was up to 75% decreased with the UCP1K56S/K269S double mutant. The NMR-based models published for these mitochondrial carriers rose serious concerns. It was first noted that the NMR model of UCP2 obtained from NMR experiment on the purified protein in DPC was instable in lipid bilayer and permeable to water molecules. In addition, the native UCP1 protein purified in DPC from brown adipose tissue was also irreversibly inactivated showing that DPC is not a suitable detergent to study UCP1 activity.^23^ In this report, we challenge the UCP1 fatty acid binding site identified by Zhao and colleagues in a mitochondrial context.^22^ We expressed UCP in yeast mitochondria and assayed uncoupling of respiration from ATP synthesis directly on permeabilized spheroplasts by oxygen consumption measurement. Wild type and UCP1 mutants were tested for their FFA mediated mitochondrial uncoupling activity.

## Results and Discussion

Yeast expression system has been extensively used either to study UCP1 function in mitochondria or permeabilized spheroplasts or to purify the protein for biophysical studies.^24–29^ Here, we designed a new construct containing an Histidin-Tag at the N-terminal of the rat UCP1 followed by a TEV cleavage site and inserted the corresponding cDNA into pYEDP60 yeast expression vector.^30^ Mutations identified in Zhao *et al*.^22^ as crucial for the proton transport activity of UCP1 are K56S, K269S, K56S/K269S. They were introduced in pYEDP60-UCP1 construct as well as the control mutation R54S.^22^ Figure 1A shows immunodetection of the recombinant UCP1 and of yeast VDAC from yeast mitochondrial extracts. All four mutants were correctly targeted to the mitochondria. Immunodetection of UCP1 was performed in triplicate on the same blot (Figure 1B-C) and statistical analysis of the variance (Supplementary Tables 1-2) confirmed that all UCP1 mutants were expressed at levels similar to wild type UCP1. Next, oxygen consumption was measured on permeabilized spheroplasts as previously described.^31^ Expression of rat UCP1 in yeast decreased the respiratory control ratio (RCR) to 1.39 +/-0.03 SEM (supplementary Figure 1) suggesting a basal proton conductance of UCP1 in yeast mitochondria as previously observed for rodent UCP1 but not for the human form.^32^ In HEK293 human cells, mouse UCP1 exhibited no basal proton conductance^33^, which can be explained either by a lower levels and of either endogneous FFA or of reactive oxygen species necessary for UCP1 activity.^34^ All UCP1 mutants K56S, K269S, K56S/K269S exhibited RCR similar to the wild type, showing that those mutations are without effect on the basal uncoupling activity of UCP1. To probe the FFA dependent proton conductance of UCP1, we investigated the uncoupling effect of fatty acids on wild type and on UCP1 recombinant yeast spheroplasts. In the absence of lauric acid, the basal level of respiration measured upon addition of NADH and oligomycin was 75% higher in spheroplasts expressing UCP1 as compared to control spheroplasts. Increasing concentration of lauric acid (60 µM to 240 µM) in presence of 15 µM free-fatty acid BSA were tested. At (LA)_60µM_/BSA_15µM_=4, respiration increased of 44% in UCP1 containing mitochondria. At (LA)_120µM_/BSA_15µM_=8 mitochondrial respiration increased further of 35%, close to its maximal value (Figure 2). In control mitochondria, addition of LA at (LA)_120µM_/BSA_15µM_=8 had no significant effect but further addition of LA stimulated respiration to its maximal value also. Therefore BSA/LA ratio of 4 and 8 were selected to assess the activation of all mutants by lauric acid. Figure 3 shows, example of respiration curves of recombinant yeast spheroplasts harboring either pYeDP, pYeDP-UCP1 or pYeDP-UCP1(K56S/K269S) expression plasmid. In the later (Figure 3A), respiration is stimulated by adding lauric acid at LA_60µM_/BSA_15µM_=4 and LA_60µM_/BSA_15µM_=8 and fully inhibited by ATP. Addition of lauric acid at LA_60µM_/BSA_15µM_=4 had no effect on control spheroplasts (Figure 3B). To confirm this finding, respiration on spheroplasts harboring either pYeDP-UCP1(K56S/K269S), pYeDP-UCP1(K56S), pYeDP-UCP1(K269S) or pYeDP-UCP1(R54S), a control mutation described in^22^, were recorded in biological triplicates. None one of them exhibited significant impaired proton transport activity (Figure 4 and Supplementary Tables 3-4 for statistical analyses). UCP1 phenotype in yeast mitochondria resembles to the physiological state of a brown adipocyte mitochondria. On whole spheroplasts, the presence of UCP1 impacts the respiratory control ratio as it has been observed in brown adipose tissue mitochondrial from many rodent species^35,36^ and from *Echinops telfairi* a protoendothermic eutherian mammal.^37^ In the yeast cell environment, UCP1 uncoupling activity is highly sensitive to low concentration of fatty acids (30 µM lauric acid with a LA/BSA=4) and fully inhibited by purine nucleotides. Therefore, the mutants that were identified based on the UCP1 NMR model and on NMR titration of UCP1 in DPC with fatty acids, do not alter the physiological uncoupling of mitochondrial respiration from ATP synthesis. Our results reinforce previous studies on UCP1 mutants produced in yeast. Garlid and colleagues^38^ found no alteration of UCP1(K269Q) activity and regulation for this mutant produced in yeast mitochondrial and reconstituted in liposomes. Gonzalez-Barroso *et al*.^*27*^ found similar results with UCP1(K269L) mutant: a proton conductance stimulated by fatty acids and inhibited by purine nucleotides. Mitochondrial carriers have been produced at high levels in *E. coli* but they form inclusion bodies, which require refolding procedure with harsh detergent. The first refolding protocol was published by the group of Palmieri and consisted of Sarkosyl detergent solubilization of inclusion bodies followed by rapid incorporation in liposomes. By doing so, they were able to establish the transport activity of the oxoglutarate carrier^39^ and of many other members of the yeast mitochondrial carrier family^40^. However this procedure is not compatible with structural analyses especially by NMR. In contrast, DPC is a commonly used detergent in NMR experiment and up to 80% of published structures of membrane proteins by NMR have been obtained in DPC.^41^ We recently reviewed the use of DPC in functional and structural biology of membrane proteins.^41^ DPC solubilizes membrane proteins at very high yield. However there are multiple examples of complete loss of function when MP are maintained in solution by DPC even at low concentration. A global analysis showed destabilizing and denaturing properties of this class of detergent especially for fragile alpha–helical membrane proteins, which mitochondrial carriers belong to.^41^ Indeed, two correspondences^42,43^ were published objecting the structural relevance of mitochondrial ADP/ATP transporter NMR based model described in Brüschweiler *et al*.^44^ Both correspondences demonstrated that AAC in DPC is fully unfolded and that the dynamic observed upon nucleotide binding is mediated by DPC detergent and has therefore no functional relevance. In response to these objections, Yang *et al*.^45^ admitted that the ADP/ATP-transporter is indeed partially folded but they argued that structural models obtained for the mitochondrial uncoupling protein 1^22^ and 2^21^, following the same biochemical approach, were functionally relevant. So far all mitochondrial carriers tested have been inactivated by DPC^23,42,43,46^ but the argument that NMR titration on a partially folded transporter would reveal crucial amino-acid for its activity persisted. Here we show that this approach is not valid for the proton transport activity of mitochondrial uncoupling protein 1. The amino-acids identified by Zhao *et al*.^*22*^ are not involved in mitochondrial uncoupling of respiration. The most simple explanation to reconciliate the results on UCP1 activity produced either in yeast or *E. coli* is that DPC interferes with UCP1 folding and activity, even after detergent removal and reconstitution of the protein in liposomes. In Zoonens *et al*., we used 60 mM DPC to solubilize brown adipose tissue mitochondria and purify UCP1 on hydroxyapatite column. At this DPC concentration, native UCP1 is irreversibly inactivated.^23^ In Zhao *et al*.^*22*^, NMR titration was also performed at 60 mM DPC concentration but the authors claimed to have partially restored the activity of the wild type UCP1 by decreasing the amount of DPC to 6 mM prior to incorporation in liposomes. Although this approach has been successful to restored the oligomeric state of the vp7 protein^47^, one possibility to explain the loss of activity of UCP1(K269S) and UCP1(K56S/K269S) is that those proteins are more fragile than the wild type and are thus irreversibly inactivated by DPC. Of note, the amount of protein incorporated in liposomes for each mutant is not documented in Zhao *et al*. and variations of the amount of protein present in liposomes would simply explain the apparent activity of each mutant. The stabilization hypothesis is supported by the fact that cardiolipin have been shown to stabilize UCP1^48^, and that structural models deriving from AAC crystal structure suggest that cardiolipin bind in the vicinity of UCP1(K56) and UCP1(K269).^17,18^ It is therefore possible that mutation K269S weakened binding of cardiolipin to UCP1, which in turn, decreased the stability of this UCP1 mutant upon reconstitution in liposomes. Our results also illustrate that, even in yeast mitochondria, the FFA concentration windows in which the specific mitochondrial uncoupling activity of UCP1 is detectable is very narrow. At (LA)_120µM_/BSA_15µM_=8 mitochondrial respiration is already stimulated in yeast harboring the control expression vector pYeDP60. The protonophoric effect of FA on mitochondria is largely documented on mitochondria, due to its genuine interaction with membrane proteins, including the ADP/ATP carrier^49^, and due to its detergent effect at high concentration. Indeed, FFA also increases the proton permeability of synthetic liposomes at concentration as low as 50 µM lauric acid.^50^ In Zhao *et al*. the effect of palmitic acid on control liposomes is not presented and it cannot be excluded that it represent a significant part of the proton leak observed in UCP1-loaded liposomes. The identification of free fatty acids binding site in UCP1 remains a difficult challenge for biochemists and is mandatory to understand how, at molecular level, proton transport occurs. Our results show that UCP1(K269) and UCP1(R56) are not involved in UCP1 proton transport activity of the protein, which highlight the difficulty to use DPC in structure-function studies of mitochondrial carriers.

**Figure 1.**
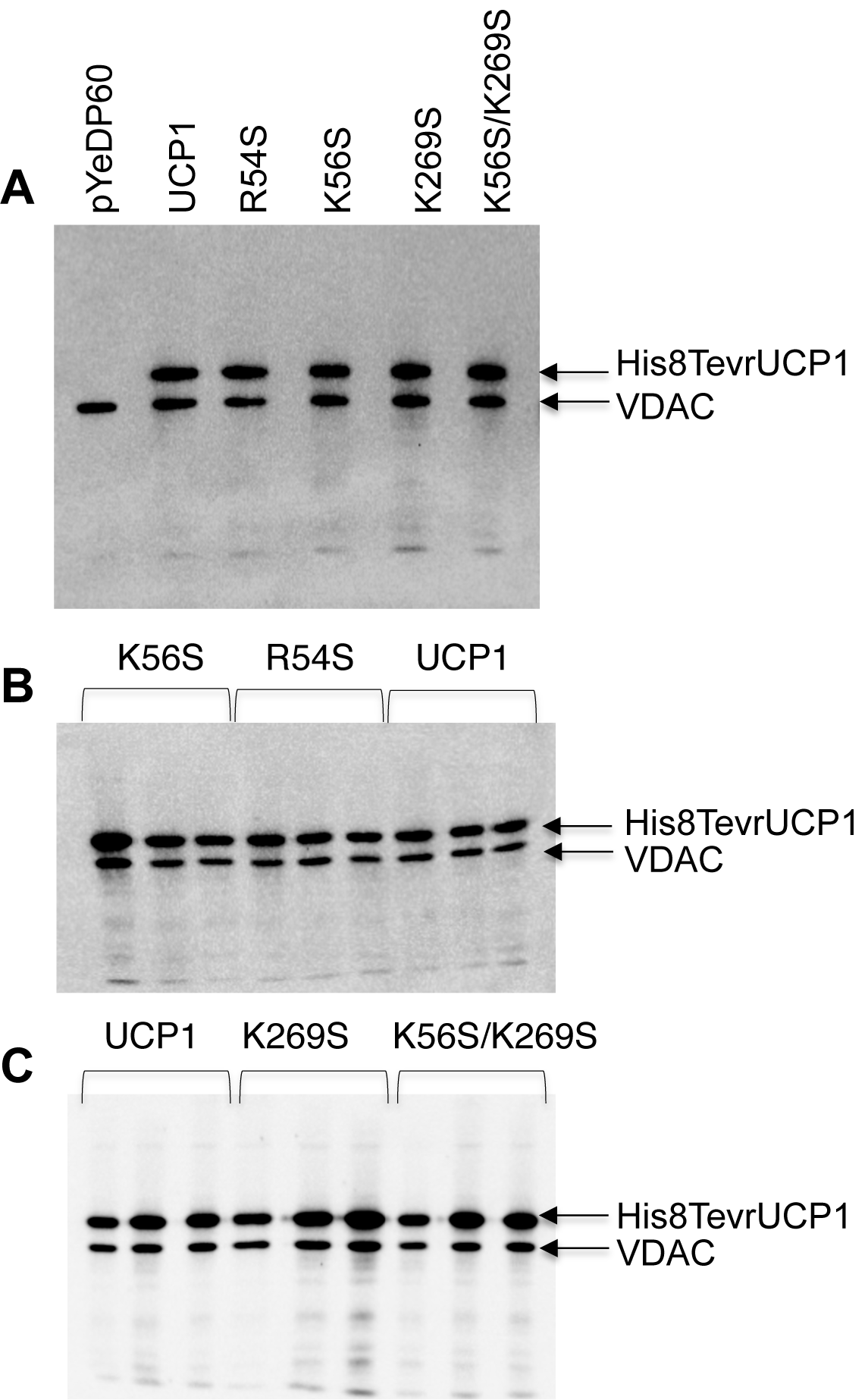
Immunodetection of UCP1 and UCP1 mutants on recombinant yeast mitochondria. Mitochondria were prepared from yeast spheroplasts and UCP1 was revealed using a HisProbe-HRP conjugate together with VDAC as control (see online methods). A. Mitochondrial extract of all UCP1 mutants versus pYeDP60. B and C. Triplicate analysis for each mutant. See supplementary Table 1 for variance analysis.

**Figure 2.**
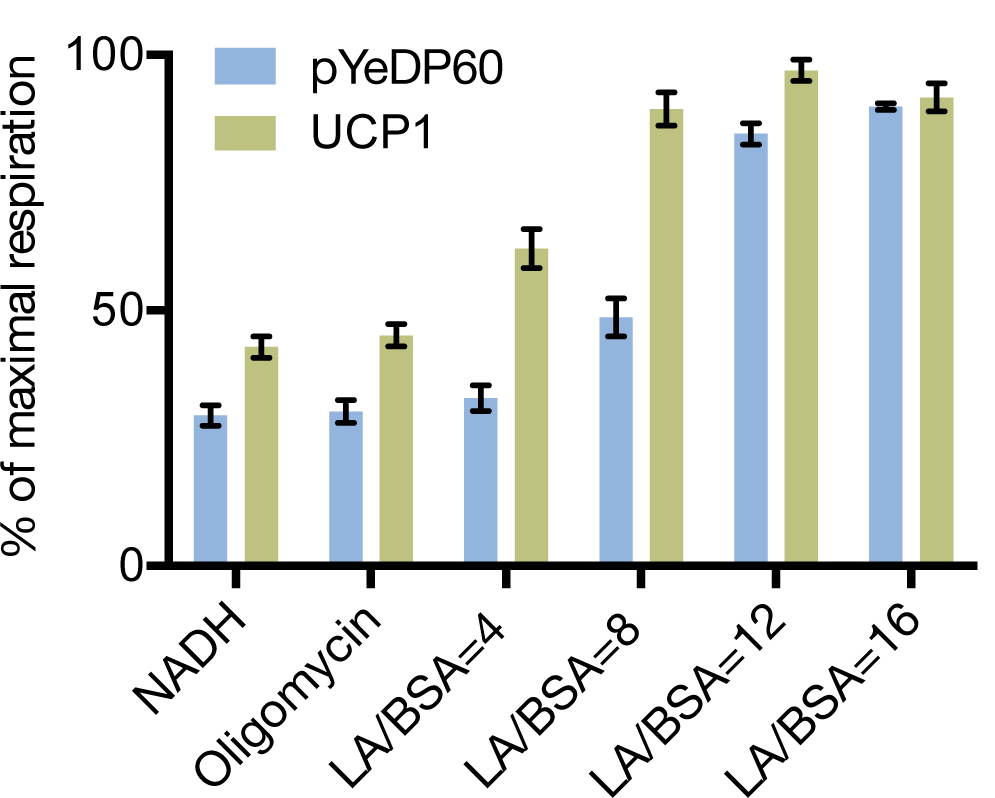
Stimulation of mitochondrial respiration by lauric acid on permeabilized spheroplasts harboring pYEDP60 and pYEDP60-UCP1 vectors. Spheroplasts were prepared according to ^31^ and permeabilized with nystatin 60 μg.ml^-1^ (see online methods). Briefly, measures were done at 25°C or 28°C in buffer containing 1 M sorbitol, 0.5 mM EDTA, 2 mM MgSO_4_, 1.7 mM NaCl, 0.1% BSA (15 μM) and 10 mM KPO_4_ pH 6.8. Oxygen consumption of permeablized spheroplasts (0.1-0.4 mg/ml) was assayed using the Oroboros instrument. NADH (3.125 mM) and oligomycin (0.5 μM) were added to induce state 4 respiration. Fatty acid mediated proton leak was induced by sequential addition of lauric acid (60 μM LA for LA/BSA=4). At the end of the experiment CCCP (5 μM) was added to calculate the maximal respiration. The graph represents the mean O_2_ flux, expressed as percentage of maximal mitochondrial respiration, upon addition of increasing concentration of lauric acid. See supplementary information for statistical analysis of variance.

**Figure 3.**
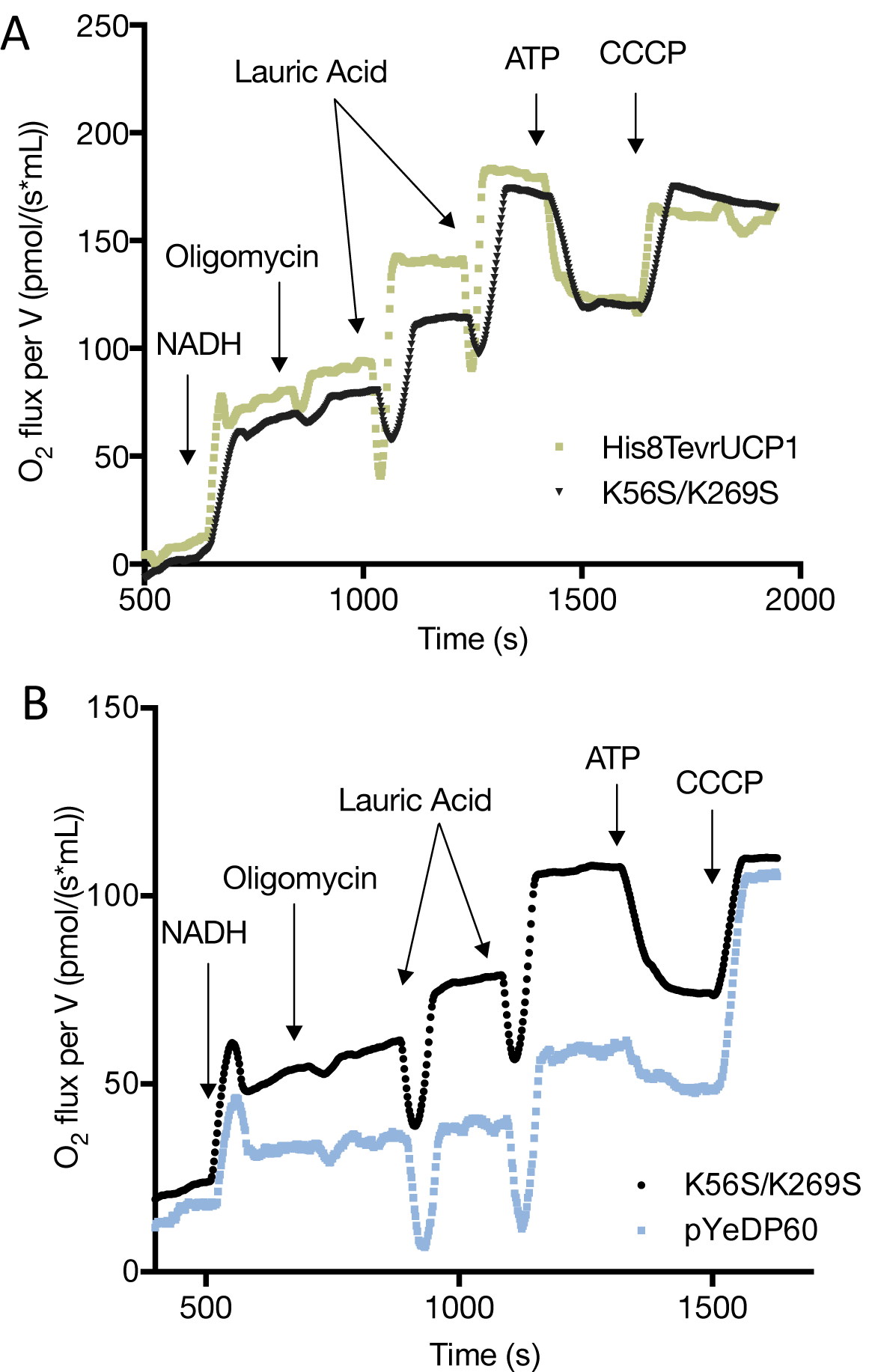
Oxygen consumption from permeabilized spheroplasts. Spheroplasts harboring either wild type UCP1, UCP1K56S/K269S mutant or pYEDP60 control expression vector were prepared as described in ^31^. Experiments were performed as described in Figure 2. Inhibition of UCP1 was achieved by addition of ATP (0.625 mM). Each graph represents a typical experiment on the Oroboros instrument loaded with two independent spheroplast samples. A, wild type UCP1 (green curve) and UCP1K56S/K269S (black curve) mutant. B, UCP1K56S/K269S (black curve) mutant and control pYEDP60 expression vector (blue curve).

**Figure 4.**
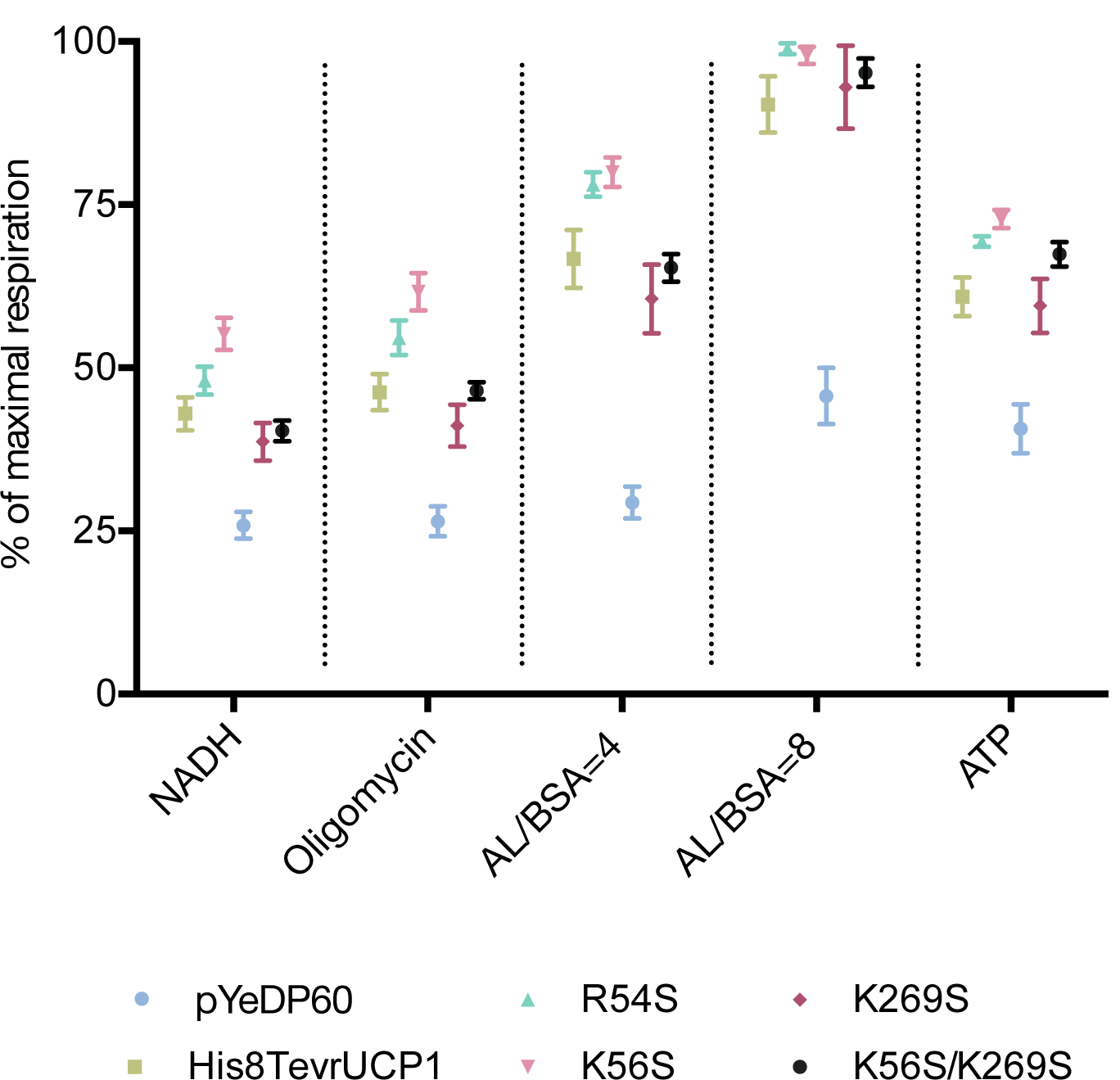
Graphical representation of mean O_2_ flux calculated from respiration curves for UCP1 and all UCP1 mutants. Experiments were performed with at least three biological replicates and one to four technical replicates. Error bars represent SEM (see supplementary information for statistical analysis of variance).

## ‘Declarations’

### Ethics approval and consent to participate

Not applicable

### Consent for publication

Not applicable

### Authors’ contributions

All authors designed the experiments and analyzed the data. MP, SM, and KM performed the experiments and analyzed the data. KM and BM wrote the paper with contributions from all authors and are co-last authors and co-corresponding authors.

## Acknowledgments

This work was supported by the Centre National de la Recherche Scientifique, and by the “Initiative d’Excellence” program from the French State (Grant “DYNAMO", ANR-11-LABEX-0011-01). MP is supported by a Ministry of Research and Technology fellowship, and BM by INSERM.

