## Supplementary data for "Structural models of mitochondrial uncoupling proteins obtained in DPC micelles are not physiologically relevant for their uncoupling activity"

Uncoupling protein 1, mitochondria, brown adipose tissue, fatty acid binding site

### Supplementary data:

1. **Expression level of R54S, K56S, and wild type UCP1 in yeast mitochondria.**

**Supplementary Table 1.**

| Mitochondrial preparation[1] | Intensity ratio[2] UCP1/VDAC | Mean ratio UCP1/VDAC | Standard Deviation[3] | Standard Error of Mean[3] |
|---|---|---|---|---|
| $WT_1$ | 1.49 | | | |
| $WT_2$ | 1.37 | 1.40 | 0.08 | 0.05 |
| $WT_3$ | 1.33 | | | |
| $R54S_1$ | 1.51 | | | |
| $R54S_2$ | 1.44 | 1.44 | 0.06 | 0.04 |
| $R54S_3$ | 1.38 | | | |
| $K56S_1$ | 1.52 | | | |
| $K56S_2$ | 1.50 | 1.61 | 0.18 | 0.10 |
| $K56S_3$ | 1.82 | | | |

[1] Mitochondria were prepared from three independent cultures; [2]Ratio of the mean intensity values were calculated with the software Image Lab 5.2.1 from the blot presented in Figure 1B; [3]Calculated with PRISM software;

Differences between mutants were not significant (P values >0.05) based on a One way ANOVA and Tukey's multiple comparisons test done on PRISM.

2. **Expression level of K269S, K56S/K269S and wild UCP1 in yeast mitochondria.**

**Supplementary Table 2.** Quantification of recombinant UCP1 and UCP1K269S and UCP1 K56S/K269S mutant expression levels in yeast mitochondria.

| Mitochondrial preparation[1] | Intensity ratio[2] UCP1/VDAC | Mean ratio UCP1/VDAC | Standard deviation[3] | Standard error of mean[3] |
|---|---|---|---|---|
| WT | 1.49 | | | |
| WT | 1.90 | 1.67 | 0.21 | 0.12 |
| WT | 1.63 | | | |
| K269S | 1.55 | | | |
| K269S | 1.89 | 1.73 | 0.17 | 0.10 |
| K269S | 1.76 | | | |
| K56S/K269S | 1.81 | | | |
| K56S/K269S | 2.11 | 2.15 | 0.36 | 0.21 |
| K56S/K269S | 2.54 | | | |

[1] Mitochondria were prepared from three independent cultures; [2]Ratio of the mean intensity values were calculated with the software Image Lab 5.2.1 from the blot presented in Figure 1C; [3]Calculated with PRISM software;

Differences between mutants were not significant (P values >0.05) based on a One way ANOVA and Tukey's multiple comparisons test done on PRISM.

**3. Respiratory control ratio.**

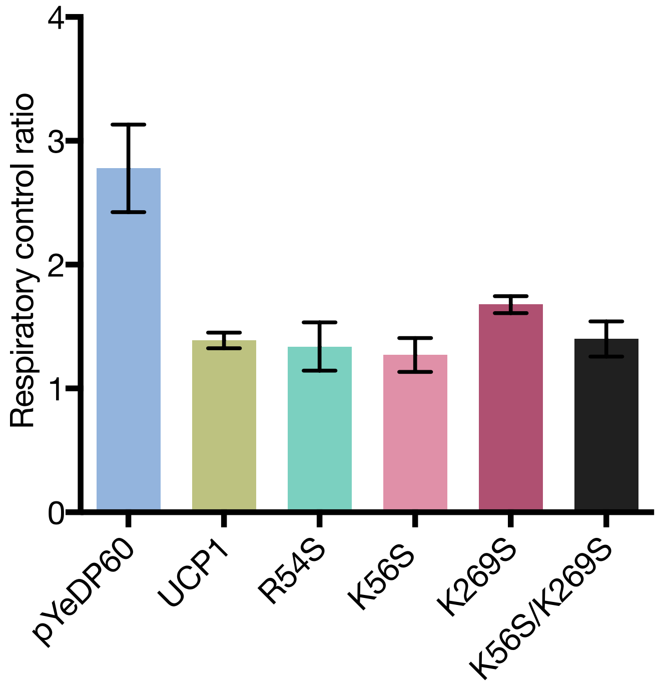


**Supplementary Figure 1.** Respiratory control ratio measured on permeabilized control spheroplasts and spheroplasts expressing mutant and wild type UCP1. Spheroplasts were permeabilized with nystatin 60 µg.ml$^{-1}$. Measures were done at 25°C or 28°C in buffer containing 1 M sorbitol, 0.5 mM EDTA, 2 mM $MgSO_4$, 1.7 mM NaCl, 0.1% BSA (15 µM) and 10 mM $KPO_4$ pH 6.8. Oxygen consumption of permeabilized spheroplasts (0.1-0.3 mg/ml) was assayed using the Oroboros instrument. Determination of the RCR was made by successive addition of NADH (3.125 mM), ADP (0.625 mM), oligomycin (0.5-1 µM) and CCCP (5 µM). The RCR was calculated by dividing the mean $O_2$ flux after addition of ADP by the mean $O_2$ flux after addition of oligomycin.

4. **Statistical analyses of spheroplast respiration with multiple t tests**

**Supplementary Table 3. Comparison between the control spheroplasts (pYeDP60) and spheroplasts expressing wild type or mutant UCP1**

| PYeDP60 vs R54S | $P_{FDR}$[1]≤0.05 | P value | pYeDP60 mean | R54S mean | Difference | SE of difference | t ratio | df[2] |
|---|---|---|---|---|---|---|---|---|
| NADH | * | 7.95E-07 | 0.26 | 0.48 | -0.22 | 0.03 | 7.35 | 18 |
| Oligomycin | * | 3.39E-07 | 0.26 | 0.55 | -0.28 | 0.04 | 7.82 | 18 |
| LA/BSA=4 | * | 3.70E-12 | 0.29 | 0.78 | -0.49 | 0.03 | 16.16 | 18 |
| LA/BSA=8 | * | 8.59E-11 | 0.46 | 0.99 | -0.53 | 0.04 | 13.38 | 18 |
| ATP | * | 1.61E-07 | 0.41 | 0.69 | -0.29 | 0.03 | 8.24 | 18 |

| pYeDP60 vs K56S | $P_{FDR}$[1]≤0.05 | P value | pYeDP60 mean | K56S mean | Difference | SE of difference | t ratio | df |
|---|---|---|---|---|---|---|---|---|
| NADH | * | 7.04E-08 | 0.26 | 0.55 | -0.29 | 0.03 | 9.00 | 17 |
| Oligomycin | * | 3.13E-08 | 0.26 | 0.62 | -0.35 | 0.04 | 9.53 | 17 |
| LA/BSA=4 | * | 2.27E-11 | 0.29 | 0.80 | -0.51 | 0.03 | 15.30 | 17 |
| LA/BSA=8 | * | 8.31E-10 | 0.46 | 0.98 | -0.52 | 0.04 | 12.15 | 17 |
| ATP | * | 1.98E-07 | 0.41 | 0.73 | -0.32 | 0.04 | 8.36 | 17 |

| pYeDP60 vs K269S | $P_{FDR}$[1]≤0.05 | P value | pYeDP60 mean | K269S mean | Difference | SE of difference | t ratio | df |
|---|---|---|---|---|---|---|---|---|
| NADH | * | 2.20E-03 | 0.26 | 0.39 | -0.13 | 0.03 | 3.69 | 15 |
| Oligomycin | * | 1.72E-03 | 0.26 | 0.41 | -0.15 | 0.04 | 3.81 | 15 |
| LA/BSA=4 | * | 4.98E-05 | 0.29 | 0.61 | -0.31 | 0.06 | 5.61 | 15 |
| LA/BSA=8 | * | 1.47E-05 | 0.46 | 0.93 | -0.47 | 0.08 | 6.28 | 15 |
| ATP | * | 4.08E-03 | 0.41 | 0.59 | -0.19 | 0.06 | 3.39 | 15 |

| pYeDP60 vs K56S/K269S | $P_{FDR}$[1]≤0.05 | P value | pYeDP60 mean | K56S/K269S mean | Difference | SE of difference | t ratio | df |
|---|---|---|---|---|---|---|---|---|
| NADH | * | 3.03E-05 | 0.26 | 0.40 | -0.14 | 0.03 | 5.62 | 17 |
| Oligomycin | * | 4.61E-07 | 0.26 | 0.47 | -0.20 | 0.03 | 7.86 | 17 |
| LA/BSA=4 | * | 2.78E-09 | 0.29 | 0.65 | -0.36 | 0.03 | 11.22 | 17 |

| | | | | | | | | |
|---|---|---|---|---|---|---|---|---|
| LA/BSA=8 | * | 6.45E-09 | 0.46 | 0.95 | -0.50 | 0.05 | 10.61 | 17 |
| ATP | * | 4.65E-06 | 0.41 | 0.67 | -0.27 | 0.04 | 6.58 | 17 |

[1] P value corrected with FDR False Discovery Rate 5 %; [2] Degree of freedom

**Supplementary Table 4. Comparison between spheroplasts expressing wild type UCP1 and spheroplasts expressing UCP1 mutants**

| UCP1 vs R54S | $P_{FDR}$[1]≤0.05 | P value | UCP1 mean | R54S mean | Difference | SE of difference | t ratio | df[2] |
|---|---|---|---|---|---|---|---|---|
| NADH | | 1.71E-01 | 0.43 | 0.48 | -0.05 | 0.04 | 1.41 | 26 |
| Oligomycin | | 4.98E-02 | 0.46 | 0.55 | -0.08 | 0.04 | 2.06 | 26 |
| LA/BSA=4 | | 5.72E-02 | 0.67 | 0.78 | -0.11 | 0.06 | 1.99 | 26 |
| LA/BSA=8 | | 1.28E-01 | 0.90 | 0.99 | -0.09 | 0.05 | 1.57 | 26 |
| ATP | | 3.44E-02 | 0.61 | 0.69 | -0.08 | 0.04 | 2.23 | 26 |

| UCP1 vs K56S | $P_{FDR}$[1]≤0.05 | P value | UCP1 mean | K56S mean | Difference | SE of difference | t ratio | df |
|---|---|---|---|---|---|---|---|---|
| NADH | * | 3.73E-03 | 0.43 | 0.55 | -0.12 | 0.04 | 3.20 | 25 |
| Oligomycin | * | 1.19E-03 | 0.46 | 0.62 | -0.15 | 0.04 | 3.66 | 25 |
| LA/BSA=4 | * | 3.82E-02 | 0.67 | 0.80 | -0.13 | 0.06 | 2.19 | 25 |
| LA/BSA=8 | | 2.03E-01 | 0.90 | 0.98 | -0.08 | 0.06 | 1.31 | 25 |
| ATP | * | 7.13E-03 | 0.61 | 0.73 | -0.12 | 0.04 | 2.93 | 25 |

| UCP1 vs K269S | $P_{FDR}$[1]≤0.05 | P value | UCP1 mean | K269S mean | Difference | SE of difference | t ratio | df |
|---|---|---|---|---|---|---|---|---|
| NADH | | 3.18E-01 | 0.43 | 0.39 | 0.04 | 0.04 | 1.02 | 23 |
| Oligomycin | | 2.75E-01 | 0.46 | 0.41 | 0.05 | 0.05 | 1.12 | 23 |
| LA/BSA=4 | | 4.17E-01 | 0.67 | 0.61 | 0.06 | 0.07 | 0.83 | 23 |
| LA/BSA=8 | | 7.31E-01 | 0.90 | 0.93 | -0.03 | 0.08 | 0.35 | 23 |
| ATP | | 7.92E-01 | 0.61 | 0.59 | 0.01 | 0.05 | 0.27 | 23 |

| UCP1 vs K56S/K269S | $P_{FDR}$[1]≤0.05 | P value | UCP1 mean | K56S/K269S mean | Difference | SE of difference | t ratio | df |
|---|---|---|---|---|---|---|---|---|
| NADH | | 4.63E-01 | 0.43 | 0.40 | 0.03 | 0.04 | 0.75 | 25 |
| Oligomycin | | 9.51E-01 | 0.46 | 0.47 | 0.00 | 0.04 | 0.06 | 25 |
| LA/BSA=4 | | 8.22E-01 | 0.67 | 0.65 | 0.01 | 0.06 | 0.23 | 25 |
| LA/BSA=8 | | 4.17E-01 | 0.90 | 0.95 | -0.05 | 0.06 | 0.82 | 25 |

| | | | | | | | |
|---|---|---|---|---|---|---|---|
| ATP | 1.30E-01 | 0.61 | 0.67 | -0.07 | 0.04 | 1.57 | 25 |

[1] P value corrected with FDR False Discovery Rate 5 %; [2] Degree of freedom

### Supplementary Materials and Methods

#### Mutagenesis:

Rat Ucp1 gene with an N terminal 8 His tag and a TEV cleavage site was cloned in pYeDP60 yeast expression vector. Site directed mutagenesis was performed using the oligonucleotides (Eurofins) presented in supplementary Table 6.

**Supplementary Table 5.**

| Mutation | Oligonucleotides |
|---|---|
| R54S | Forward: CCAGGCTTCCAGTACTATTAGTTATAAAGGTGTCTTAGGG<br>Reverse: CCCTAAGACACCTTTATAACTAATAGTACTGGAAGCCTGG |
| K56S | Forward: GCTTCCAGTACTATTAGGTATTCAGGTGTCTTAGGGACC<br>Reverse: GGTCCCTAAGACACCTGAATACCTAATAGTACTGGAAGC |
| K269S | Forward: CCGGCAGCCTTTTTCTCAGGGTTTGCGCCTTCTTTTC<br>Reverse: GAAAAGAAGGCGCAAACCCTGAGAAAAAGGCTGCCGG |

After mutagenesis subcloning was done in pYeDP60 vector. Sequences were verified by sequencing (Eurofins).

*Transformation in yeast:*

Transformation in yeast was achieved following the lithium acetate/single-stranded carrier DNA/polyethylene glycol method. *Saccharomyces cerevisiae* strain W303.gal4 was grown in YPDA medium (1% yeast extract, 2% peptone, 2% glucose, 100 mg/l adenine sulphate) overnight at 200 rpm, 30°C. At $OD_{600nm}$=7, 0.5 ml of cells were harvested by centrifugation at 5000 g for 2 min. The pellet was washed with TE buffer (10 mM Tris-HCl pH 7.5, 1 mM EDTA) and cells were pelleted and resuspended in transformation mix (500 µl PEG4000 40%, 100 mM LiOAc, 50 µl DMSO, 5 µl DNA carrier) supplemented with the pYEDP60 vectors (0.6 µg). The suspension was carefully homogenized by pipetting the mixture up to 30 times and by shaking the solution 15 min at room temperature followed by an incubation for 15 min at 42°C. Cells were then washed 3 times with TE buffer, resuspended in 150 µl of TE buffer and spread on Sdaa plate (0,67% yeast nitrogen base without amino acid, 0.5% casamino acids, 40 mg/l tryptophan, 2% agar). Plates were incubated 3 days at 30°C.

*Yeast expression and preparation of spheroplasts:*

Preculture was done overnight (30°C under 200 rpm agitation) in S-lactate medium (2% lactate, 0.67% yeast nitrogen base without amino acids, 0.1% casamino acids, 0.12%

$(NH_4)_2SO_4$, 0.1% $KH_2PO_4$, 20 mg/l tryptophan, pH 4.5) supplemented with glucose (0.1%). Preculture was diluted to $OD_{600nm}$=0.03 in S-lactate with glucose 0.1% and yeast cells were incubated 20 hours at 30°C under 200 rpm agitation to $OD_{600nm}$=3. In order to induce the expression of the recombinant protein, medium was exchanged with new S-lactate medium supplemented with 1% galactose and cells were further grown at 200 rpm 30°C for 4 hours. Cells were harvested by centrifugation, washed in water and resuspended (6 ml/g of wet yeast pellet) in 0.1 M Tris-HCl pH 9.3, 0.5 M 2-mercaptoethanol. Cells were incubated 10 min at 32°C with gentle stirring every 2 min, washed with Zymolyase buffer (1.2 M sorbitol, 20 mM $KH_2PO_4/K_2HPO_4$ pH7.4) and resuspended in 4 ml/g of yeast cells of Zymolyase buffer containing 1 mg/ml Zymolyase. After 45 min incubation at 30°C, spheroplasts were washed 3 times (800 g centrifugation, 5 min) with respiration buffer (1 M sorbitol, 0.5 mM EDTA, 2 mM $MgSO_4$, 1.7 mM NaCl, 10 mM $KH_2PO_4/K_2HPO_4$, 0.1% BSA pH 6.8) and resuspended in respiration buffer (4 ml/g of wet cells). Proteins in spheroplast preparations were quantified by BCA assay with BSA as standard. Mitochondria were isolated from spheroplasts. Spheroplasts were washed with 1.2 M sorbitol, 20 mM $KH_2PO_4/K_2HPO_4$ pH 7.4 (800 g, 5 min), resuspended in 0.6 M sorbitol, 20 mM $KH_2PO_4/K_2HPO_4$ pH 7.4 and broken by gentle homogeneization with a glass dounce potter. Mitochondria were isolated by differential centrifugation. Nuclei were harvested by centrifugation at 800 g for 5 min, and mitochondria present in the supernatant were harvested at 10 000 g for 10 min. Mitochondria were resuspended in 10 mM Tris-HCl pH 7.6, 1 mM EDTA, 250 mM Sucrose.

*Gel electrophoresis and western blots:*

Proteins in mitochondrial preparations were quantified by BCA assay with BSA as standard. Proteins (2 μg) were loaded on 11% SDS-PAGE gel and transferred onto nitrocellulose membrane for immunodetection. UCP1 was revealed using a HisProbe-HRP conjugate (Thermo Fisher Scientific 2 μg/mL) and VDAC was detected using a monoclonal anti-VDAC antibody (16G9E6BC4 Thermo Fisher Scientific 100 ng/ml in PBS tween) and an HRP-coupled goat anti mouse monoclonal antibody (Promega W402B 100 ng/ml). Detection of peroxidase activity was achieved using the Pierce ECL western blot kit (Thermo Fisher Scientific) and intensity of the bands was estimated by using Image lab software (5.2.1) with local background correction.

*Respiration experiments:*

The activity of UCP1 was assayed based on $O_2$ consumption measurement made with an Oroboros instrument at 25°C or 28°C under 750 rpm agitation. Spheroplasts (2 mg/ml) were permeabilized in respiration buffer supplemented with nystatin (66 µM) for 20 min at 28°C, diluted to 0.1-0.3 mg/ml in respiration buffer and loaded in the Oroboros recording chamber.

The respiratory control ratio (RCR) was measured by successive addition of NADH (3.125 mM), ADP (0.625 mM), oligomycin (0.5-1 µM) and CCCP (5 µM). The RCR was calculated by dividing the mean $O_2$-ADP flux by the mean O2-oligomycin flux. Respiration experiments for RCR determination were performed four times with at least two independent yeast cultures. Activity of UCP1 was revealed by adding NADH (3.125 mM), oligomicyn (0.5-1 µM) and increasing concentration of lauric acid ((LA), 60 µM to 240 µM - corresponding to a ratio LA/BSA from 4 to 16). Inhibition of UCP1 at LA/BSA=8 was obtained by adding ATP (0.625 mM). CCCP (5 to 15 µM) was added at the end of the experiment. $VO_{2max}$ was defined as the highest $VO_2$ consumption rate during one respiration experiment; it corresponded either to $VO_2$-CCCP or $VO_2$-lauric acid.

*Statistical analysis:*

Mean $O_2$ fluxes were calculated from respiration curves after each addition with a time window of 1 min. Data are presented as mean ± SEM. The statistical analysis was performed in Prism software with a multiple t test (False discovery rate =5%).
